# EVALUATION OF THE EFFECT OF MESENCHYMAL STEM CELLS ON CHEMOTHERAPY RESPONSE FOR NEUROBLASTOMA TREATMENT IN AN EXPERIMENTAL ANIMAL MODEL

**DOI:** 10.1101/2020.11.09.373936

**Authors:** Safiye Aktas, Yuksel Olgun, Hande Evin, Ayse Pinar Ercetin, Tekincan Cagri Aktas, Osman Yilmaz, Gunay Kirkim, Nur Olgun

## Abstract

High-dose cisplatin (CDDP) causes dose-limiting side effects in neuroblastoma (NB) treatment. Mesenchymal stem cells (MSC) are a current research area in cellular treatments due to multipotential characteristics. The aim of this study is to assess the interaction of MSC with CDDP in an athymic nude mouse NB model. Athymic male nude mice (n=28) were injected subcutaneously with C1300 NB cell line. After tumor growth to 1 cm diameter in 7-10 days, mice were randomly assigned to one of 4 experimental groups of control, CDDP treatment, MSC treatment and CDDP+MSC treatment with 7 mice in each group. Animals had basal auditory tests performed and had physiological serum or CDDP (20 mg/kg) injected into the peritoneum and were intravenously injected with 1×10^5^ MSC once. Seven days later, hearing tests were performed again and the animals were sacrificed. Tumor tissue was assessed in terms of necrosis, apoptosis and viability. Apoptosis was evaluated with annexin V+PI flow cytometry analysis and TUNEL. Additionally, the MSC rate within the tumor was assessed with flow cytometry for triple CD34+ CD44+ and CD117-expression. Additionally, liver, kidney, brain and cochlear tissue were analyzed with light microscopy in terms of systemic side effect profile. Expression of the cochlear cell proteins of calretinin, math-1 and myosin2A were immunohistochemically assessed in ear sections. Statistical analysis used the nonparametric Kruskal Wallis and Mann Whitney U tests with p<0.05 significance. Tumor tissues were found to have statistically significantly higher levels of necrosis in the CDDP group and CDDP+MSC group compared to the control and MSC groups (p=0.001, p=0.006). The CDDP+MSC group had lower tumor necrosis rates than the CDDP group but this was not observed to have statistical significance (p=0.05). MSC did not change the tumor dimensions in the CDDP group (p=0.557). The groups administered MSC had higher triple CD34+ CD44+ and CD117- expression within tumor tissue compared to the control and CDDP groups. In the inner ear, the expression of cochlear cell proteins calretinin, math-1 and myosin2A were identified to be highest in the groups administered MSC. Auditory tests observed that the 15-decibel loss at 12, 16, 20 and 32 kHz frequencies in both ears with CDDP was resolved with MSC administration. With this study, IV administration of MSC treatment was observed to prevent the hearing loss caused by CDDP without disrupting the antitumor effect of CDDP. Systemic MSC may be assessed for clinical use to reduce the side effects of CDDP.

## INTRODUCTION

Neuroblastoma (NB) is a noteworthy embryonal tumor with interesting heterogeneous biological behavior rooted in the neural crest of the sympathetic nervous system. In spite of intensive protocols and alternative treatment approaches in advanced stage disease, the two-year disease-free survival only reaches 30-40%. Cisplatin (CDDP) is used in the induction therapy cycle for NB in combination with anthracyclines, alkylating agents and topoisomerase II inhibitors [1]. Induction chemotherapy continues with a multimodal approach involving surgical resection, myeloablative treatment and autologous stem cell transplant and radiation therapy [2]. In recent years, immunotherapy, the ALK inhibitor crizotinib in patients with ALK mutations and targeted treatments based on genomic analysis of tumor samples have been trialed.

In NB treatment areas, side effects of hearing loss (62%), primary hypothyroidism (24%), ovarian failure (41% in women), musculoskeletal system anomalies (19%) and pulmonary anomalies (19%) are observed [5]. CDDP is used for human solid tumors in many cancer types like ovarian, prostate, cervical, head-neck, lung and bladder cancer and in NB. However, considering ototoxicity especially, nephrotoxicity, neurotoxicity and bone marrow toxicity occurring with use of high-dose CDDP, dose limitation represents a significant problem for the therapeutic profile and benefit of the drug [6,7]. Apart from ototoxicity, all other side effects may be resolved with support treatment methods. There is research into many protective agents, both chemical agents and containing natural extracts, for use against CDDP ototoxicity. The current literature includes a variety of dose- and/or time-dependent efficacy for many agents including Korean red ginseng [8], dexamethasone [9], resveratrol [10], silymarin [11], metformin [12], selenium [13] and others. In addition to preventing hearing loss occurring linked to CDDP and disrupting quality of life in a serious sense, studies about regaining this faculty have gained much importance.

Mesenchymal stem cells (MSC) have multipotent capability and have become a topic of current research as a cell transplant-based promising therapeutic approach due to their ability to differentiate into mesenchymal cell series like osteoblast, adipocyte and chondroblast. However, there are few studies interrogating the interaction of MSC with cancer treatment and preventive properties for chemotherapy side effects. MSC has natural tropism in tumor tissue. MSC transports antitumor molecules like cytokines and interferon into the tumor microenvironment and are considered as cellular treatment agents [14]. MSC were shown to mediate transformation in neurogenic differentiation of cochlear auditory hair cells in vitro [15]. Our previous studies attempted to administer MSC and CDDP to an in vitro coculture model of cochlear cells. It was concluded that MSC supported renewal of cells after ototoxicity was induced in HEI-OC1 cochlear cells by CDDP. The study induced ototoxicity with 100 uM CDDP and administered MSC with 40% difference identified in viability of cochlear cells after incubation. It was identified that MSC reduced the cochlear cell injury caused by CDDP [16]. In light of these findings, our hypothesis in this study is that systemic MSC administered with CDDP for NB will reduce the ototoxicity side effect and not change the anticancer efficacy.

The aim of the study is to assess whether systemic MSC administration in an NB experimental animal model disrupts the antitumoral effect of CDDP used as chemotherapy agent and to reveal the side effect profile, especially in terms of ototoxicity.

## MATERIALS AND METHODS

This study received research ethics committee approval from Dokuz Eylül University Multidisciplinary Animal Laboratory Animal Experiments Ethics Committee at a meeting dated 12 September 2017 with protocol number 38/2017.

### Cell Culture

#### C1300 cell line

This is a mouse-derived NB cell line. As seen in our previous studies, it is a cell line inducing tumors in athymic nude mice [17]. It was cultured in DMEM media (1% L-glutamine and 1% penicillin/streptomycin) containing 10% fetal bovine serum at 37 °C in a 5% CO2 incubator. When cells reached nearly 90% confluent levels, they were removed from the flask surface with trypsin-EDTA solution and placed in a 96-well plate with 6 wells/group and 5000 cells per well. Cells were left for 24 hours to adhere to the wells and then upper phases were removed with a pipette taking care not to lift the cells. CDDP with different doses was administered (10, 25, 50, 100 and 250 uM). The plate was incubated for 24 hours at 37 °C in a 5% CO2 incubator. At the end of the incubation duration, cell viability was examined with MTT. LD50 doses were determined [18] (Fig 1).

**Figure 1:**
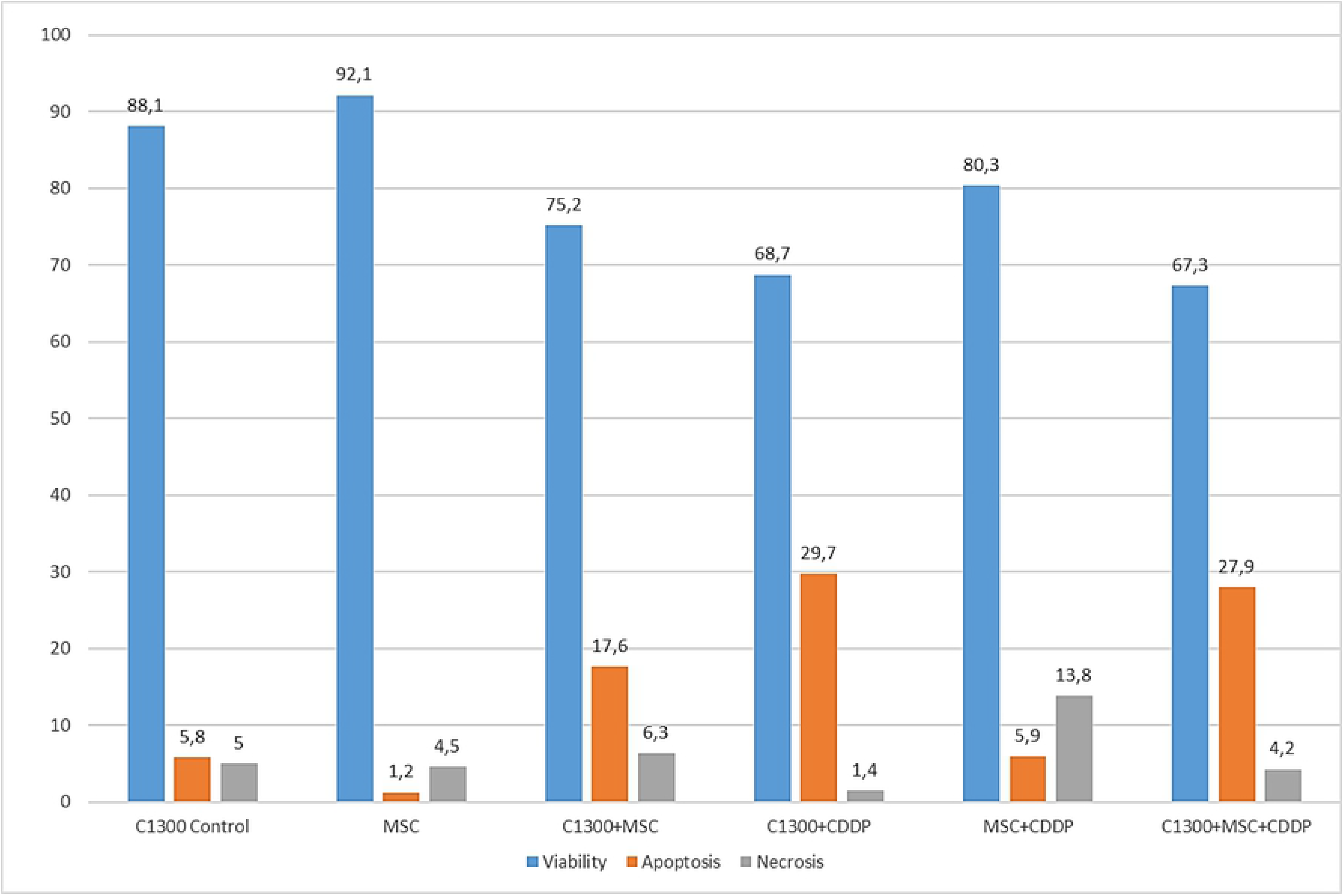
In vitro cytotoxic effects of Cisplatin on C1300 neuroblastoma cells with/ without MSCs.

#### C57BL/6 mouse mesenchymal stem cells line

This is a mesenchymal stem cell line derived from mouse bone marrow (Cyagen, `MUBMX-01201`). It has osteogenic, chondrogenic and adipogenic differentiation features. It was developed from C57BL/6 mouse tibia. It is CD34 positive, CD44 positive and CD117 negative. It was purchased at ninth passage. It was cultured in a cell-specific differentiation preventing medium at 37 °C in 5% CO2 humidified incubator (OriCellTMMouseMesenchymal Stem Cell Growth Medium (Cat. No.MUXMX-90011)). Medium was changed once every two days to reduce MSC markers and it was dissociated and passaged with trypsin-EDTA when 75% confluent. Freezing was performed in a protein-free cryopreservation freezing medium.

#### Coculture formation

To assess the effect of MSC cells on the cytotoxic effect of CDDP for NB cells, C1300 cells were proliferated in a 96-well plate. With 6 wells for every set, the control group only had medium applied, the CDDP group had LD50, the only MSC group had 5000 MSC, the CDDP and MSC group had CDDP LD50 + 5000 MSC applied for 24 hours. Cell viability was tested with MTT. Under the same conditions, apoptosis cell death was assessed with annexin V+PI flow cytometry in 6 wells.

### Cell Viability Test

Cell proliferation testing with MTT used 96-well plates containing cells, with only medium placed in three empty wells used blind. With cells of 100 μl/well in each well, 10 μl/well MTT cell proliferation reactive was added (1:10 dilution). Cells were left at 37 °C 5% CO2 in an incubator for 4 hours. After mixing on a plate mixer for 1 min, they were read with an ELISA reader at 420-480 nm. The reference wavelength was chosen as more than 600 nm (630 nm). Mean absorbance in the control group was accepted as 100% viability, while this was compared with other cell viability levels to obtain percentages.

### Induction of Xenograft Neuroblastoma Tumor Model

In our study, male athymic nude mice (nude CD1 mice) aged 8 weeks with mean weight 20 g were placed in a special room in Dokuz Eylül University Faculty of Medicine Experimental Animals Research Laboratory (DEÜTFDHAL). Ventilation used a hepafilter, and mice were housed in special cages at room temperature (20 ± 2 °C) with 12-hour light/dark environment. They were fed with sterile pellet mouse feed and given access to sterile water ad libitum. Before beginning the study, mice were observed in this environment for one week to ensure they adapted. Study design and time schedule is given in fig 2.

**Figure 2:**
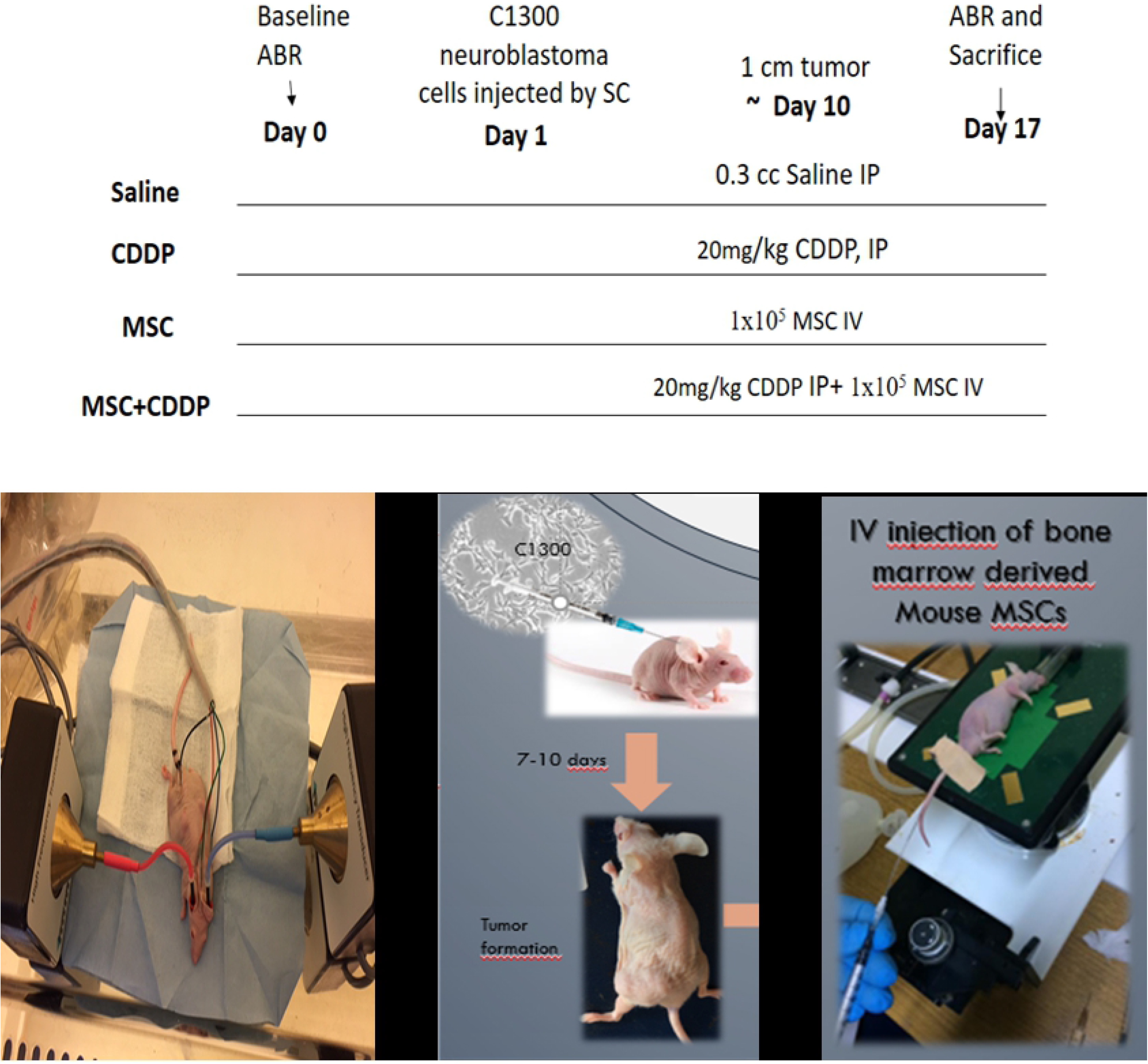
Study design and time shedule.

Mice had C1300 NB cells with 1×10^6^ cell/0.3 ml in incomplete DMEM injected via the subcutaneous route on the left side of the back to ensure tumor development. When the induced tumors reached 150 mm3 (nearly 10 days), mice were randomized into groups with n=7 in each four group (Control SF, MSC, CDDP, MSC+CDDP). Number of mice per group (sample size) was selected prioritizing 3R rule and using our previous studies in order to achieve statistical power greater than 80% at 0.05 alpha by ClinCalc program sample size calculator. Independent study groups with dichotomous primary end point for hearing loss was taken for anticipated incidence 15% in control group and 85% in CDDP group. If no tumor formation was observed within 15 days of this administration, the same amount of C1300 cells were injected again to assess tumor formation. During the study, the plan was that animals with development of sepsis, with veterinary agreement and with clear reduction in response to stimuli, and with loss of more than 15% weight would be excluded from the study. There was no need to exclude any animals from the study. Mice were weighed routinely once per day in a class 2 cabinet to monitor weight loss. The researchers performed all studies in sterile conditions wearing gloves, glasses and special aprons. Tumor size was monitored each day with calipers. One week after CDDP (20 mg/kg) and MSC (106 cell/mouse) administration, mice were sacrificed (fig 3). After applications, remaining medication solutions were appropriately destroyed. Before sacrificing mice, isoflurane inhalation anesthesia was administered. The abdomen was opened and nearly all blood was removed from the vena cava inferior and the cardiac main veins were cut. After death, organs and tissues of mice were dissected. After dissecting the residual tumor bed, part of it was stored in a cell culture medium, while part of it was placed in formoline solution for light microscope analyses.

**Figure 3:**
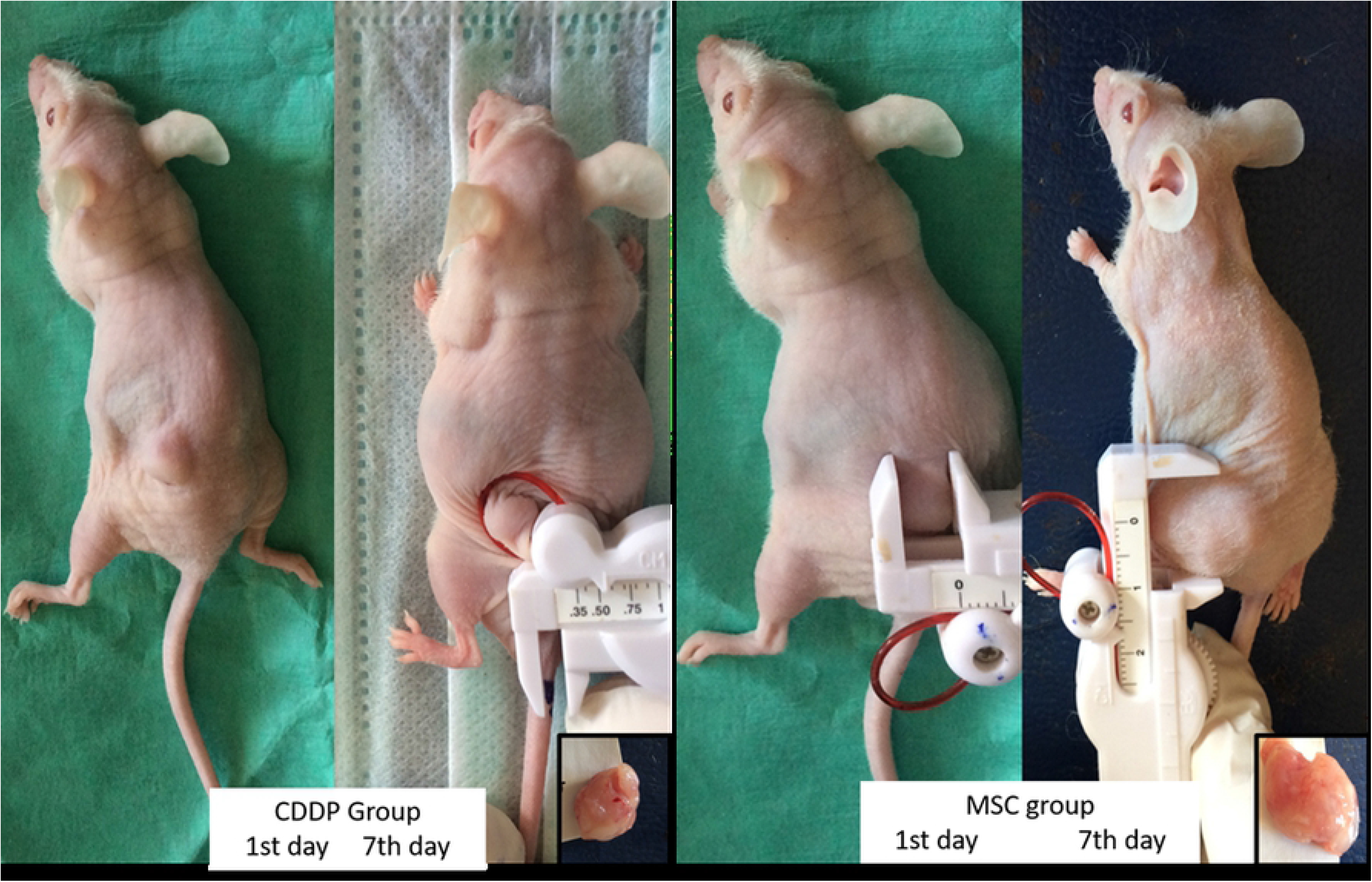
Subcutanous Tumor appearances of nüde mice comparing first and seventh days in CDDP and MSC groups.

#### Chemicals

##### Cisplatin (KOÇAK FARMA)

Liquid form was administered at dose of 20 mg/kg under in vivo conditions to mice with cancer induced.

### Brainstem Auditory Evoked Potential Test

Nude mice were administered anesthesia with 40 mg/kg ketamine and 10 mg/kg xylazine intraperitoneal and basal and 7^th^ day evoked brainstem responses were tested with the ABR test (Fig 4). All nude mice first had otoscopic examination performed before hearing measurements and nude mice with normal otoscopic exam and basal ABR test hearing threshold of 25 dB SPL and below were included in the research. Tests were completed under appropriate aseptic conditions and in sterile cabinets. An Intelligent Hearing Systems (IHS, Miami, FL) device Smart-EP 10 version was used. Subdermal needle electrodes were inserted into the vertex, ipsilateral and contralateral retroauricular areas. A platinum-iridium needle electrode was used as recording electrode. The ABR test used 4, 8, 12, 16, 20 and 32 kHz in the Blackman envelope with tone burst stimuli with fluctuation time 1000 ms [19, 20]. The lowest intensity level obtained with the III wave was accepted as the hearing threshold of the nude mouse at that frequency.

**Figure 4:**
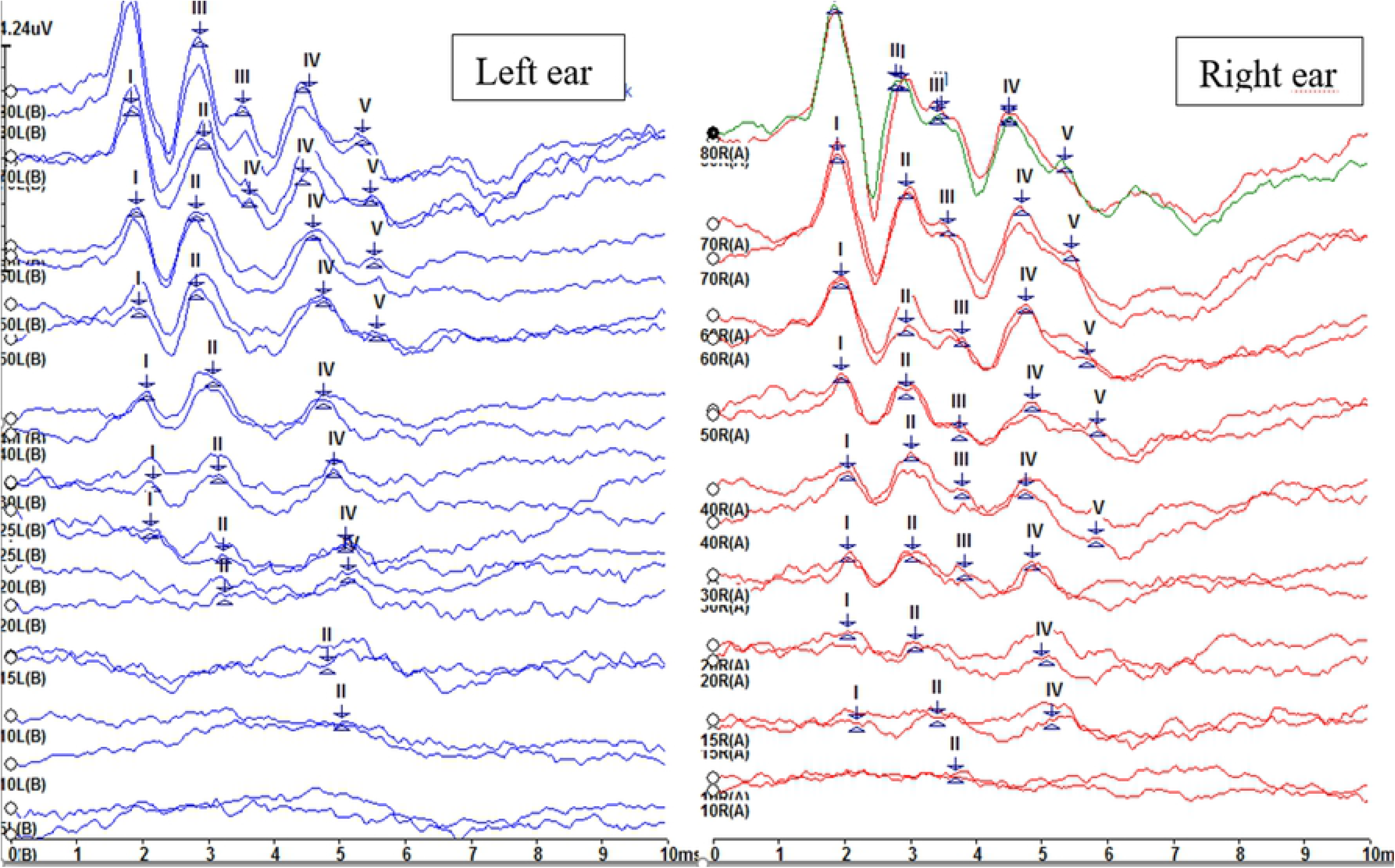
ABR records of right and left ear.

### Light Microscopic Assessment

Half of tumor tissue, kidney, lung, brain, cerebellum, heart, spleen and liver were left in formol for 24 hours after dissection. Sections were chosen with macroscopic investigations and cut to cassettes. The region including the outer, central and inner ear involving bone tissue was dissected from the skull by an ear-nose-throat expert. After fixation in formol for 24 hours, it was left in 5% glacial acetic acid for 3 days for decalcification. It was cut in two from the broad surface and left in acid for one more day, washed with flowing water and cut to sections with 3 mm thickness as cassettes. All cassettes underwent standard large tissue monitoring and paraffin blocking after being left in formol for 24 hours and then sections with 5-micrometer thickness were placed on slides and stained with hematoxylin eosin. These slides were examined histopathologically by a pathologist under a light microscope (Olympus B50). Tumor tissue was researched for morphology, angiogenesis, invasion, necrosis and live tissue proportions and organs were examined for morphological changes.

### Apoptosis Analysis with TUNEL on Paraffin Sections

Sections obtained after routine paraffin blocking processes for half of tumor tissue and ear tissue sections were left overnight in a 60 °C incubator and then left in 2 × 30 min xylene after cooling. Sections were passed thought 96%, 80%, 70% and 60% ethyl alcohol series for 2 minutes each and washed with PBS for 5 min. Circumference of sections were drawn with a pen and left in Proteinase K solution at 1:500 dilution for 15 min at room temperature. After 3 × 5 min washing with PBS solution, endogenous peroxide blockage was performed (3% H2O2) (5 min). After washing with PBS solution for 3 × 5 min, they were washed in equilibration buffer solution for 5 min at room temperature. For each section, 100 μl TdT solution was prepared (77 μl reaction buffer solution + 33 μl TdT) and dropped onto the sections. Sections with TdT had a plastic lamella placed on the slide and were left at 37 °C for 1 hour and then washed at room temperature for 10 min with prepared reaction stopping buffer solutions (1 ml stop washing buffer + 34 ml distilled water). They were left with anti-digoxygenin conjugate for 30 min at room temperature. After washing with PBS solution for 3 × 5 min, DAB solution was dropped on the sections and they were left in an enclosed humid box for 5-10 min. After washing with PBS solution for 3 × 5 min, they were washed with distilled water and nucleus staining was performed with Mayer’s hematoxylin checked with 1-5 min staining. After washing with distilled water, the sections were left in 80%, 96% and 100% ethyl alcohol for 1 min each, then dried and purified with xylene 2 times for 5 min each. Closing medium was used and sections were covered with lamella. Assessment counted 1000 cells in 5 different areas of the tumor tissue and recorded the mean percentages. In the inner ear, all cells and positive cells were counted and % values were calculated.

### Apoptosis Assessment with In Vitro Tests and Fresh Tumor Tissue Annexin V+PI

Cells collected with a cell scraper during in vitro tests and created by mechanical degradation of half of the tumor tissue from the 28 animals when fresh were passed through a 50-micrometer filter and stored at −80 °C in single cell suspension freezing medium. Half of the suspended cells were placed in 15 ml Falcon tubes, centrifuged at 1200 rpm for 5 min, supernatant removed and then resuspended with 1 × concentration 100 μl buffer prepared by dilution of the 10 x binding buffer given with the pellet kit and transferred to polystyrene 5 ml flow cytometry test tubes. In order to perform correct gating settings for the analysis, the same sample was placed in one tube without staining, while another tube had only 5 μl propidium iodide (PI), one tube had only 5 μl FTIC-Annexin V and one tube had PI with Annexin V added to 5 μl. These tubes were incubated for 15 min at room temperature in a dark environment. At the end of incubation, the tube had 400 μl binding buffer added and flow cytometry analysis was performed. After appropriate gating, those stained with only Annexin V were identified as early apoptotic, those stained with both annexin V and PI were late apoptotic and those with only <<< PI staining were identified as necrotic. The percentage data obtained were used for comparative calculations according to the initial cell counts.

### Determination of Intratumoral Mesenchymal Stem Cell Counts

The other half of the single cell suspension created by mechanically degrading half of the fresh tumor tissue after sacrifice in medium and passing through a 50-micrometer filter was used. CD34 (abcam PerCP Cy 5.5), CD44 positive (abcam APC), and CD117 negative (abcam PE) MSC markers were assessed with flow cytometric analysis. Flow cytometric analysis of MSC biomarkers was completed using monoclonal antibodies labeled with PerCP Cy 5.5, PE and APC after incubation at room temperature for 15 min in a dark environment and then washing with PBS. After centrifuging, unbound antibodies were removed and the sample was resuspended in 500 μl PBS and analyzed with a BD Accuri flow cytometry device. MSCs identified as CD34+ CD44+ CD117-were identified with appropriate gating.

### Assessment of Mesenchymal Stem Cell Differentiation in the Inner Ear

After fixation and decalcification, serial sections were cut from paraffin blocks with a microtome and sections with 5 micrometer thickness from the inner ear region were placed on positive-loaded slides. Unstained serial sections were investigated under the microscope and sections containing Corti organ and spiral ganglion were chosen and separated for immunohistochemical staining. To show the stimulating effect of MSC on cochlear cell differentiation preventing the ototoxic effect of CDDP on cochlear cells, specific markers for MSC and HEI-OC1 in the experimental groups were identified with immunocytochemical staining. MSC markers used CD34 (abcam), while HEI-OC1 biomarkers used myosin IIA (Bioss), Math-1 (Bioss), and calretinin (Bioss) staining. Immunohistochemical staining was performed with automatic staining in a Ventana Discover device. The device deparaffinizes sections and places them in water. Cells are made permeable for intracellular staining and then blocking, incubation with 1/200 diluted primary antibodies, incubation with secondary antibodies and appropriate washing steps are performed. Then contrast staining with hematoxylin is performed, sections are dehydrated by treatment with xylol, closed with a lamella and evaluated with a light microscope. Calretinin is a marker of immature inner ear as expression begins in the inner ear from the 13^th^ day of the embryonic period and continues until the adult period. Math-1 is expressed by developing hair cells and is not expressed by non-sensorial cells. Math-1 shows positive expression in immature and hair cells [21–23].

### Statistical Analysis

SPSS 22.0 software was used to compare means for viability, apoptosis and differentiation markers in every experimental group (non-parametric Mann Whitney U test and Kruskal Wallis test) including data from 7 animals in every group. Comparison of categoric variables was performed with the chi-square Fisher exact test. P<0.05 was accepted as statistically significant.

## RESULTS

### In Vitro Results

The viability percentages of C1300 cells after 24 hours with 25 uM, 50 uM, 100 uM, 250 uM and 500 uM CDDP treatment were 88.1%, 68.7%, 51.3%, 43.7%, and 32.5%, respectively; while after 48 hours with 25 uM, 50 uM, 100 uM, 250 uM and 500 uM CDDP treatment viability was 42.9%, 34.2%, 22.6%, 16.2% and 11.2%, respectively. The viability percentages for C1300 cells after 72 hours of 25 uM, 50 uM, 100 uM, 250 uM and 500 uM CDDP treatment were 92.3%, 91.3%, 77.8%, 38.5% and 19.2%, respectively. Accordingly, for in vitro tumor modelling, 100uM doses of CDDP and 24 hours of incubation were selected. After MSCs co-cultured with C1300 cells were treated with 25 uM, 50 uM, 100 uM, 250 uM and 500 uM of CDDP for 24 hours, cell viability percentages were 61%, 70%, 48.8%, 31.1% and 17.5%, respectively. When C1300 cells and MSCs were cocultured and treated with CDDP, the selected time and dose did not change.

### Xenograft Neuroblastoma Tumor Model Results

Mice with mean tumor diameter 1.2 mm had medication, cell and physiological saline administered. Difference in tumor diameters from initial values over 7 days were noted in terms of tumor progression. The difference in mean tumor diameter was 4.85 mm in the control group, 2.14 mm in the MSC group, −3.14 mm in the CDDP group and −2.43 mm in the CDDP+MSC group. While the tumor diameters increased in the control and MSC groups, the tumor diameters reduced in the CDDP and CDDP+MSC groups. Both CDDP and CDDP+MSC administrations reduced tumor diameter compared to the control group (p=0.01). Administration of MSC alone was not identified to cause a statistical difference in tumor diameter difference compared with the control group (p=0.128). Administration of MSC did not statistically significantly change the reduction in tumor size compared to the CDDP group (p=0.805).

### Apoptosis Assessment with Annexin V+PI in Fresh Tumor Tissue

Single cell suspension was prepared from tumor tissue on the 7^th^ day and analyzed with flow cytometry. When the control group is compared with the CDDP group, higher early apoptotic, late apoptotic, total apoptotic and necrotic cell percentages were identified in the CDDP group (p=0.017, 0.038, 0.026, 0.001). Again, when the control group is compared with the CDDP+MSC group, cell measurements for all four parameters were found to be higher in the CDDP+MSC group (p=0.001). Both apoptotic and necrotic cell death was observed in the CDDP and CDDP+MSC groups. When administered with MSC, cell death did not change statistically (p=0.165, 0.62, 0.318, 0.535, respectively). When the control group and MSC groups are compared, no differences were identified for the four parameters (p=0.62, 0.62, 0.805, 0.71). Systemic MSC administration did not affect tumor cell death.

Mean apoptosis and necrosis percentages are given in Table 1.

**Table 1:** Mean Percentages of Flow Cytometry Annexin V+PI Cell Death Parameters

| Group | % | Early Apoptosis | Late Apoptosis | Total Apoptosis | Necrosis |
| --- | --- | --- | --- | --- | --- |
| Control |  | 0.8 | 6.34 | 7.14 | 3.98 |
| CDDP |  | **14.07** | **15.14** | **29.21** | **21.03** |
| MSC |  | 1.66 | 5.57 | 7.23 | 6.21 |
| MSC+CDDP |  | **13.57** | **15.29** | **28.86** | **20.16** |

### Light Microscopic Assessment

Mean necrosis of tumor tissues was 12.85% in the control group, 57.14% in the CDDP group, 3.71% in the MSC group and 35.00% in the MSC+CDDP group. Hematoxylin eosin staining of tumor tissues found necrosis was higher in the CDDP group and CDDP+MSC group compared to the control and MSC groups (p=0.001, p=0.011). The CDDP+MSC group appeared to have lower tumor necrosis rates compared to the CDDP group (p=0.017). Tumor sections observed solid tumor growth patterns in live tumor areas in all four groups. There was undifferentiated appearance. Differentiation findings were not observed in rosette formation. Vascularization was pronounced. Both pushing growth in the form of nodules and infiltrative growth was observed in subdermal fat connections and muscle tissue. Postmortem investigations of mice found no areas of suspected metastasis of macroscopic tumors. Notable pathology was not observed in other organs. Histopathology of liver, lung, kidney, heart, spleen and brain tissue sections was normal.

There was no degradation in histomorphology of inner ear sections. There was no flat epithelium (FE) appearance secondary to cellular damage in severe sensorial hearing loss. The cytoarchitecture of the Corti organ was normal. Interior and exterior hair cells could be selected. However, there were pyknotic changes supporting cell death in the spiral ganglion of the CDDP group.FE is seen in humans and mice with profound sensorineural hearing loss and/or vertigo. Various factors, including ototoxic drugs, noise exposure, aging, and genetic defects, can induce FE. Both hair cells and supporting cells are severely damaged in FE, and the normal cytoarchitecture of the sensory epithelium is replaced by a monolayer of very thin, flat cells with irregular contours.

### Determination of Mesenchymal Stem Cell Counts with Flow Cytometry in Tumor Tissue

In the control group and the CDDP group, triple CD34+, CD44+ and CD117-expression was not identified, while it was 3% in the MSC group and 5.16% in the MSC+CDDP group. Flow cytometry of single cell suspension from tumor tissue assessed with triple labeling identified higher MSC numbers in tumor tissue of groups administered MSC compared to the control and CDDP groups, as expected (p=0.001). There was no statistical difference between the MSC and MSC+CDDP groups. These findings show systemic administration of MSC entered tumor tissue through circulation [21].

Expression of cochlear cell proteins calretinin, math-1 and myosin2A in the inner ear were identified to be higher in the group administered MSC. On hearing tests, the 15-decibel loss at 12, 16, 20 and 32 kHz frequencies with CDDP in both ears was observed to be resolved with administration of MSC [22–24].

### Apoptosis Analysis with TUNEL on Paraffin Sections

Mean apoptotic cells in tumor tissue were identified as 2.2% (2-8) in the control group; 30% (5-45) in the CDDP group; 5.43% (1-15) in the MSC group; and 28.57% (15-55) in the CDDP+MSC group. Statistical analysis identified more apoptosis in the CDDP and CDDP+MSC groups compared to the control and MSC groups (p=0.007, 0.001, 0.004). There was no difference between the CDDP and CDDP+MSC groups (p=0.456) (Fig 5) (Fig 6).

**Figure 5.**
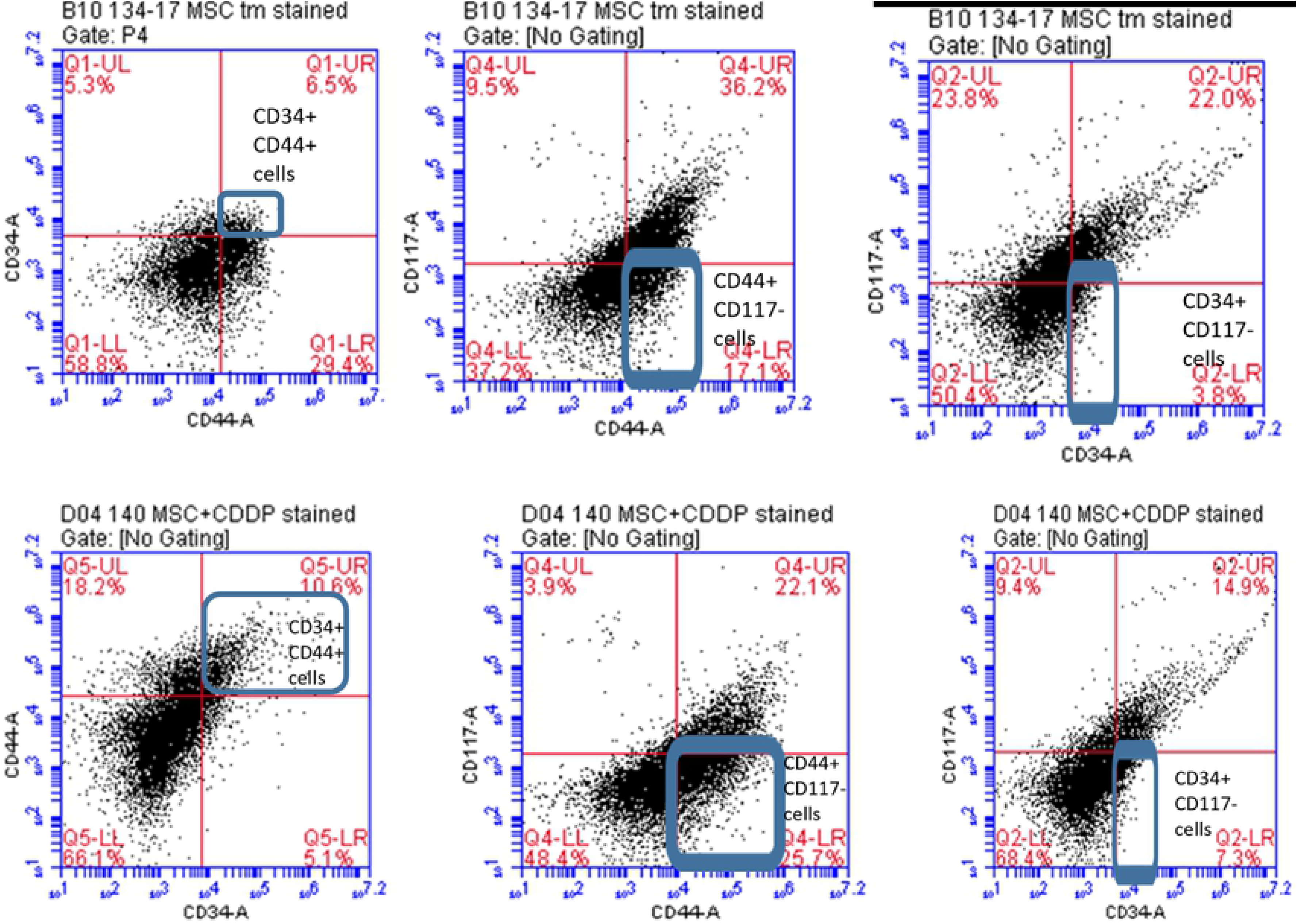
MSC (CD34+ CD44+ and CD117-) flow cytometer of of fresh tumor tissue single cell suspension in IV MSC applied (upper 3 figs) and MSC+CDDP applied (lower 3 figs) mice.

**Fig 6:**
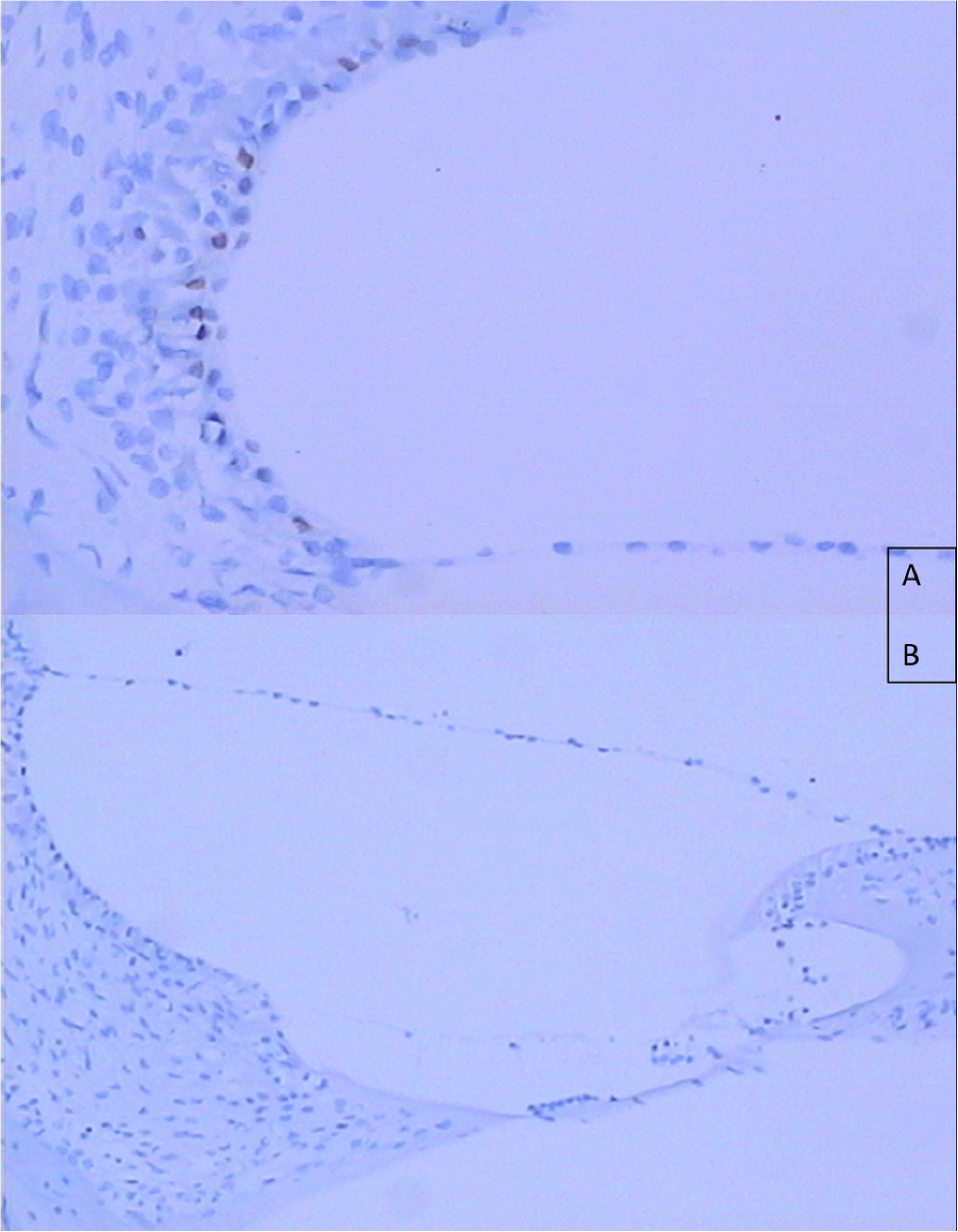
Apoptotic cells stained in brown color in Stria vascularis layer of cohlea in CDDP group (TUNEL method, DAB, x400) (A). No apoptosis was observed in Control group (B) (x200)

Apoptosis was not identified in the Corti organ in the inner ear, while values in the spiral ganglion were mean 26% and 28% in the CDDP and CDDP+MSC groups. Statistically significant high levels of apoptotic cells were identified in these groups compared to the control and MSC groups (p=0.001).

### Brainstem Auditory Evoked Potential Test Results

There was no statistically significant difference between groups in terms of baseline ABR values (p>0.05). According to ABR measurements conducted on the 7th day of the study, hearing thresholds for all frequencies except 8 kHz were significantly deteriorated in the CDDP-treated group in comparison to the other 3 groups (p<0.05). In the CDDP + MSC treated group, hearing thresholds were not statistically different from controls and the MSC treated group (Fig 7).

**Figure 7.**
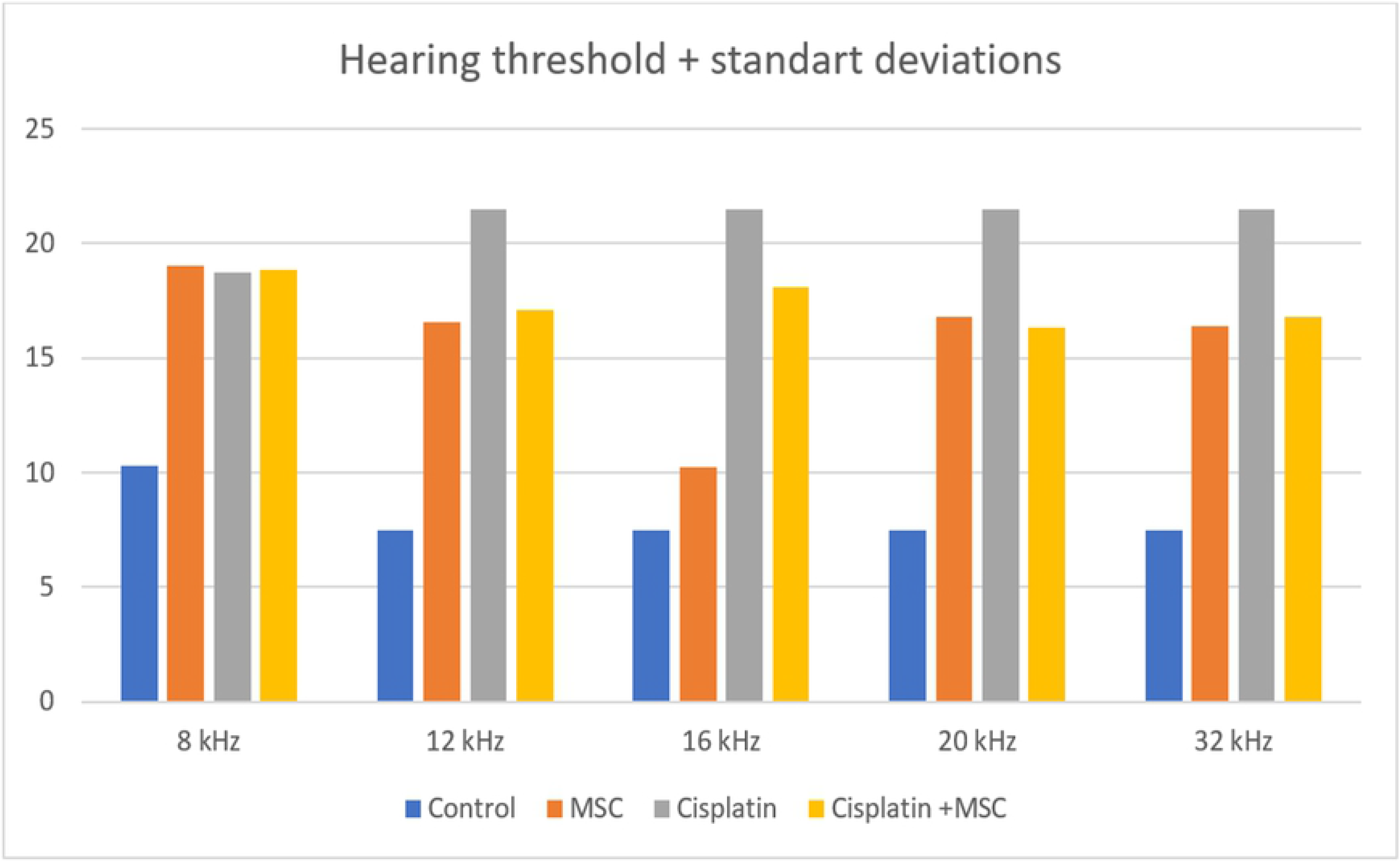
Mean hearing thresholds + standart deviations in all groups at 7th day of the study.

### Assessment of Mesenchymal Stem Cell Differentiation in the Inner Ear

Calretinin begins to be expressed in the inner ear on the 13^th^ day of the embryonic period and is a marker that continues to be expressed until the adult period. In the control, CDDP and MSC+CDDP groups, calretinin expression was positive in the organ Corti and spiral ganglion. In the MSC group, calretinin expression was negative in organ Corti tissues. Systemic administration of MSC reduced the expression of calretinin in ear tissue.

Developing hair cells express Math-1, while it is not expressed by non-sensorial cells. Math-1 has positive expression in immature and hair cells. In the control and CDDP groups, low expression of Math-1 was identified. In the MSC group and MSC+CDDP groups, Math-1 expression was increased (p=0.04).

Myosin2A plays an active role in reorganization within the cochlea. Suppression of this antigen causes defects in the cellular pattern. Myosin2A had positive expression pattern in all control, CDDP, MSC and MSC+CDDP groups. There was no difference observed between the groups (p>0.05).

Among these three parameters, Math-1 expression increased in the inner ear after systemic administration of MSC and it undertakes a marker role for cochlear effect.

## DISCUSSION

Sensorineural hearing loss caused by CDDP of 60% in the childhood period and 50% in adults disrupts quality of life. As a result, development studies for potential protective agents are currently in progress [25]. However, apart from the use of sodium thiosulfate in childhood cancers, there is no agent with proven reliability and benefit at the level of clinical studies. Agents used in current pre-clinical trials include CDK2 inhibitors, G-protein coupled receptor (GPCR) agonists and lovastatin. In our study, we obtained findings that systemic MSC cellular treatment simultaneous to CDDP reduced the ototoxicity side effect without disrupting the antitumoral effect at the level of in vivo animal experiment studies. We assessed this effect in terms of apoptotic and necrotic effects. MSC was tested with triple markers, and we attempted to measure the contribution to the protective mechanism against damage with immature cochlear cell markers in the inner ear.

A limitation of our study is that we did not include a local administration model for transtympanic, intracochlear MSC. We could not model MSC cell administration within the inner ear at the dimension of nude mice under our laboratory conditions. Our study only includes systemic intravenous MSC administration through the tail vein. It is very difficult for many medications to access the interior of the cochlea. CDDP induces hearing loss by damaging sensorial cells especially external hair cells, stria vascularis cells and spiral ganglion cells in the inner ear [26]. Migration of MSC to this region via circulation is not an expected situation. We think the effect is possibly due to mediation by MSC-derived soluble factors in an environment with systemic oxidative stress.

Kasagi et al. labeled MSC with green fluorescent protein in culture medium and transplanted them into the posterior semicircular canal of mice. At the end of this study, MSC had differentiated to fibrocyte-like cells in the cochlea and did not harm hearing functions [26]. Lin et al. researched an inductive method to differentiate MSC derived from bone marrow into hair cells in the cochlea. As a result of culturing MSC in medium without serum containing EGF, IGF-1 and N2/B27 in coculture with neurons for two weeks, they observed differentiation into cells like hair cells expressing myosin VIIa [27]. If embryonic stem cells locally administered to the inner ear lower high potassium concentration within toxic endolymph, they showed that they may survive in normal ear epithelium and flat epithelium for up to 7 days. However, there is a need for new studies about hair cell differentiation [27]. In our study we investigated the effects of systemic MSC.

Lopez-Juarez et al. stimulated pluripotent stem cells to obtain otic progenitor cells and they transplanted these cells into the cochlea in an ototoxic guinea pig model. These cells remained viable for 4 weeks and showed molecular features of early sensorial differentiation [28]. This current study opened the horizons in terms of treatment of ototoxicity with stem cells. However, there are important gaps in relation to organization of cells in the organ Corti. Our study is different to this study in terms of examining prevention of ototoxicity with systemic MSC administered simultaneous to CDDP chemotherapy, which causes hearing loss without severe cellular injury,

## Conclusion

For NB treatment, protective agents are frequently researched to reduce side effects due to the dose-limiting effects of CDDP side effects. It is essential that these agents do not disrupt the anti-tumor effect. In our study, systemic administration of MSC as cellular treatment was identified to have a protective effect against the ototoxicity side effect of CDDP and did not disrupt anti-tumoral efficacy in a nude mice NB model. Planning of clinical studies may be considered; however, firstly it is necessary to investigate the effect mechanisms of MSC on the organ Corti in terms of differentiation, regeneration, systemic effect and epithelial mesenchymal transformation.

## Acknowledgement

This study was presented as a poster numbered 8-1238 at the International Society of Pediatric Oncology (SIOP) Congress in 2018. This project was financially supported by Dokuz Eylül University Rectorate Scientific Research Projects Coordinator with project number 2018.KB.SAG.076 for purchasing of consumables.

